# An improved mode of running PASTA

**DOI:** 10.1101/2020.08.30.274217

**Authors:** Qikai Yang, Tandy Warnow

## Abstract

PASTA is a method for estimating alignments and trees that has been able to provide excellent accuracy on large sequence datasets. By design, PASTA operates using iteration, in which the tree from the previous iteration is used to inform a divide-and-conquer strategy during which a new alignment is computed on the sequence dataset, and then a new maximum likelihood tree is estimated on the new alignment. In its default setting, PASTA runs for three iterations and returns that alignment/tree pair from the last iteration. Here we use both biological and simulated nucleotide datasets to show that returning the alignment/tree pair that has the best maximum likelihood score improves on the default usage.

## 1 Introduction

Multiple sequence alignment (MSA) is a fundamental step in many biological studies, and large-scale MSA is particularly difficult. Several methods have been designed for large-scale MSA, including Clustal-Omega (Sievers et al., 2011), KAlign3 (Lassmann, 2019), PASTA (Mirarab et al., 2015), and UPP (Nguyen et al., 2015). Furthermore, sequence length heterogeneity also presents challenges for alignment (Smirnov and Warnow, 2020), a problem that is addressed by UPP. Specifically, when given a dataset with sequence length heterogeneity, UPP extracts a set of sequences that are “full length” and uses PASTA to align them, thus forming the “backbone alignment”. The remaining sequences are then added into the backbone alignment using an ensemble of Hidden Markov Models (Durbin et al., 1998). Thus, UPP is an extension of PASTA to enable the accurate estimation of alignments given sequence length heterogeneity. Hence, here we focus on PASTA, due to its high accuracy on large datasets with high rates of evolution and applicability to both nucleotide alignment and protein alignment.

PASTA uses a combination of divide-and-conquer and iteration, so that each iteration operates by taking the alignment and tree from the previous iteration, dividing the sequence input into subsets using the tree, aligning the subsets using the selected base method, and then merging the alignments together. A maximum likelihood tree is then estimated on the alignment using FastTree2 (Price et al., 2010). This process repeats for several iterations (default 3), and then the last alignment/tree pair is returned. While this default usage has performed well in many studies, here we explore the potential for additional improvement in accuracy by returning the alignment/tree pair that had the best maximum likelihood score, as computed using FastTree2.

## 2 Methods and materials

### 2.1 Methods

We compare two variants of PASTA, the default version and a variant where we return the alignment/tree pair with the best maximum likelihood score, on a collection of biological and simulated datasets with up to 1000 sequences, each run for four iterations, and we report the impact on alignment error. All datasets are available in public repositories from prior studies.

PASTA-BestML differs from the default settings for PASTA in two ways: it does not mask sites that are highly gappy (as is performed in PASTA-Default, in order to speed up the analysis on large datasets) and it returns the alignment/tree pair that has the best maximum likelihood score across all the reported iterations (instead of the last alignment/tree pair). Since PASTA has some randomness in its execution, we ran several independent experiments on each dataset. Therefore, we ran 100 independent experiments for each biological dataset and 50 independent experiments for each replicate of each simulated dataset.

### 2.2 Datasets

We include both biological nucleotide datasets and simulated nucleotide datasets from prior studies. We use a standard protocol to preprocess these datasets to reduce to a set of sequences that are full-length: we discard all sequences whose length is different from the median sequence length by over 20%. We use two sources for simulated datasets: the 1000- and 500-sequence datasets used in Liu et al. (2009, 2011) to evaluate SATé and SATé-II in comparison to other methods, which were simulated using ROSE (Stoye et al., 1998), and some subsets of the RNASim dataset (studied in Mirarab et al. (2015)) with 1000 sequences; the ROSE datasets are available at Mirarab (2020) and the RNASim datasets are available at Smirnov (2020).

All the simulated datasets evolve with substitutions and indels, but differ from each other in various ways. The ROSE datasets from Mirarab (2020) evolve under a modification of the GTRGAMMA model to allow for indels, and the RNASim datasets evolve with substitutions and indels under a biophysical model that reflects the selective pressure of RNA structure conservation (see Mirarab et al. (2015)). The ROSE datasets include a wider range of sequence evolution and thus include some conditions that are challenging to align, while the RNASim datasets are also challenging due to highly variable rates of evolution across the length of the sequences. Each of the simulated datasets has 20 replicates. We include six biological nucleotide datasets from the Comparative Ribosomal Website (CRW) (Cannone et al., 2002), available from Mirarab (2020). They are 16S.M, 16S.M.aa_ag, 23S.M, 23S.M.aa_ag, 23S.E and 23S.E.aa_ag; these range in size from 96 to 740 sequences, and from 930 nucleotides to 3599 nucleotides in average sequence length. See Tables 1 and 2 for empirical statistics about these datasets.

**Table 1:** Information on biological nucleotide datasets from Mirarab (2020), and originally from Cannone et al. (2002). The p-distance of a pair of aligned sequences was calculated by dividing the number of sites where the two sequences had different nucleotides by the number of sites in which both sequences had nucleotides.

| Datasets | Average<br>p-distance | Average<br>seq. length | # sequences<br>(after screening) | # sequences<br>(before screening) |
| --- | --- | --- | --- | --- |
| 16S.M | 0.298 | 947 | 740 | 901 |
| 16S.M.aa_ag | 0.300 | 930 | 633 | 1028 |
| 23S.E | 0.283 | 3599 | 99 | 117 |
| 23S.E.aa_ag | 0.284 | 3531 | 96 | 144 |
| 23S.M | 0.326 | 1580 | 211 | 278 |
| 23S.M.aa_ag | 0.339 | 1578 | 197 | 263 |

**Table 2:** Information on simulated nucleotide ROSE datasets from Mirarab (2020) and RNASim datasets from Smirnov (2020).

| Datasets | Average<br>p-distance | Average<br>seq. length | Average<br># sequences<br>(after screening) | Average<br># sequences<br>(before screening) |
| --- | --- | --- | --- | --- |
| ROSE-1000L1 | 0.695 | 1015 | 1000 | 1000 |
| ROSE-1000L2 | 0.696 | 1007 | 1000 | 1000 |
| ROSE-1000L3 | 0.687 | 1031 | 1000 | 1000 |
| ROSE-1000L4 | 0.500 | 1007 | 1000 | 1000 |
| ROSE-1000L5 | 0.496 | 1006 | 1000 | 1000 |
| ROSE-1000M1 | 0.695 | 1011 | 1000 | 1000 |
| ROSE-1000M2 | 0.684 | 1014 | 1000 | 1000 |
| ROSE-1000M3 | 0.660 | 1007 | 1000 | 1000 |
| ROSE-1000M4 | 0.495 | 1006 | 1000 | 1000 |
| ROSE-1000M5 | 0.499 | 1003 | 1000 | 1000 |
| ROSE-1000S1 | 0.694 | 1002 | 1000 | 1000 |
| ROSE-1000S2 | 0.693 | 1001 | 1000 | 1000 |
| ROSE-1000S3 | 0.686 | 1002 | 1000 | 1000 |
| ROSE-1000S4 | 0.501 | 1000 | 1000 | 1000 |
| ROSE-1000S5 | 0.498 | 1000 | 1000 | 1000 |
| ROSE-500L1 | 0.670 | 1042 | 500 | 500 |
| ROSE-500L2 | 0.657 | 1037 | 500 | 500 |
| ROSE-500L3 | 0.658 | 1023 | 500 | 500 |
| ROSE-500L4 | 0.499 | 1023 | 500 | 500 |
| ROSE-500L5 | 0.497 | 1010 | 500 | 500 |
| ROSE-500M1 | 0.674 | 1017 | 500 | 500 |
| ROSE-500M2 | 0.658 | 1018 | 500 | 500 |
| ROSE-500M3 | 0.657 | 1009 | 500 | 500 |
| ROSE-500M4 | 0.491 | 1011 | 500 | 500 |
| ROSE-500M5 | 0.495 | 1004 | 500 | 500 |
| ROSE-500S1 | 0.673 | 1004 | 500 | 500 |
| ROSE-500S2 | 0.655 | 1004 | 500 | 500 |
| ROSE-500S3 | 0.656 | 1002 | 500 | 500 |
| ROSE-500S4 | 0.492 | 1002 | 500 | 500 |
| ROSE-500S5 | 0.498 | 1000 | 500 | 500 |
| RNASim-1000 | 0.411 | 1555 | 1000 | 1000 |
| RNASim-1000_C_100 | 0.317 | 1551 | 1000 | 1000 |
| RNASim-1000_C_500 | 0.378 | 1555 | 1000 | 1000 |

### 2.3 Criteria

We report alignment error using SPFP and SPFN, calculated by FASTSP (Mirarab and Warnow, 2011), where SPFN is the percentage of the pairwise homologies in the reference alignment that do not appear in the estimated alignment and SPFP represents the percentage of the homologies in the estimated alignment that are missing from the reference alignment.

### 2.4 Dataset Information

## 3 Results and Discussion

As seen in Table 3 and Table 4, PASTA-BestML has lower alignment error than PASTA-Default for all biological datasets and nearly all simulated datasets. Thus, running PASTA with BestML typically improves alignment accuracy compared to default PASTA. The difference is generally small, but because it is nearly universal, it suggests that PASTA-BestML may be a more reliable technique for using PASTA than its default mode. This improvement also suggests that the maximum likelihood score may be a valuable criterion to use when selecting an alignment from a set of candidate alignments.

**Table 3:** Comparison between PASTA’s default mode and PASTA’s BestML mode for alignment error (SPFP/SPFN) on biological nucleotide datasets. For each dataset and criterion, the best result is boldfaced.

|  | SPFP |  | SPFN |  |
| --- | --- | --- | --- | --- |
|  | Default | BestML | Default | BestML |
| 16S.M | 14.22% | **14.01%** | 12.54% | **12.31%** |
| 16S.M.aa_ag | 13.20% | **13.17%** | 11.46% | **11.43%** |
| 23S.E | 21.04% | **20.61%** | 17.03% | **16.58%** |
| 23S.E.aa_ag | 20.39% | **20.14%** | 16.41% | **16.12%** |
| 23S.M | 22.72% | **22.67%** | 17.80% | **17.71%** |
| 23S.M.aa_ag | 24.12% | **24.06%** | 18.70% | **18.64%** |

**Table 4:** Comparison between PASTA’s default mode and PASTA’s BestML mode for alignment error (using SPFP/SPFN) on simulated nucleotide datasets, taken from Mirarab (2020) (simulated using ROSE) and Smirnov (2020) (simulated using RNASim). For each model condition and criterion, the lower error rate is boldfaced at each row.

|  | SPFP |  | SPFN |  |
| --- | --- | --- | --- | --- |
|  | Default | BestML | Default | BestML |
| ROSE-1000L1 | 8.48% | **7.54%** | 8.68% | **7.75%** |
| ROSE-1000L2 | 3.22% | **3.08%** | 3.29% | **3.14%** |
| ROSE-1000L3 | 16.36% | **15.66%** | 16.84% | **16.15%** |
| ROSE-1000L4 | 0.60% | **0.59%** | 0.60% | **0.59%** |
| ROSE-1000L5 | **0.39%** | 0.40% | **0.39%** | 0.40% |
| ROSE-1000M1 | 18.18% | **17.28%** | 18.85% | **17.95%** |
| ROSE-1000M2 | 13.98% | **13.40%** | 14.34% | **13.76%** |
| ROSE-1000M3 | 4.81% | **4.67%** | 4.82% | **4.69%** |
| ROSE-1000M4 | **1.01%** | 1.02% | **1.00%** | 1.01% |
| ROSE-1000M5 | 0.60% | **0.59%** | 0.60% | **0.59%** |
| ROSE-1000S1 | 14.99% | **13.91%** | 15.32% | **14.25%** |
| ROSE-1000S2 | 7.52% | **6.83%** | 7.58% | **6.90%** |
| ROSE-1000S3 | 5.82% | **5.45%** | 5.85% | **5.48%** |
| ROSE-1000S4 | 0.54% | **0.53%** | 0.54% | **0.53%** |
| ROSE-1000S5 | 0.21% | **0.20%** | 0.21% | **0.20%** |
| ROSE-500L1 | 18.07% | **17.64%** | 18.71% | **18.29%** |
| ROSE-500L2 | 14.70% | **14.40%** | 15.04% | **14.73%** |
| ROSE-500L3 | 6.77% | **6.59%** | 6.79% | **6.60%** |
| ROSE-500L4 | 1.67% | **1.65%** | 1.64% | **1.62%** |
| ROSE-500L5 | 0.74% | **0.73%** | 0.72% | **0.71%** |
| ROSE-500M1 | 17.02% | **16.52%** | 17.73% | **17.18%** |
| ROSE-500M2 | 12.27% | **12.05%** | 12.63% | **12.41%** |
| ROSE-500M3 | 6.20% | **6.14%** | 6.27% | **6.22%** |
| ROSE-500M4 | 1.38% | **1.36%** | 1.37% | **1.35%** |
| ROSE-500M5 | 0.76% | **0.73%** | 0.75% | **0.73%** |
| ROSE-500S1 | 15.89% | **15.28%** | 16.26% | **15.62%** |
| ROSE-500S2 | 11.05% | **10.74%** | 11.21% | **10.88%** |
| ROSE-500S3 | 5.28% | **5.07%** | 5.30% | **5.09%** |
| ROSE-500S4 | 1.70% | **1.69%** | 1.68% | **1.67%** |
| ROSE-500S5 | 0.74% | **0.73%** | **0.73%** | **0.73%** |
| RNASim-1000 | 10.08% | **9.90%** | 9.97% | **9.78%** |
| RNASim-1000_C_100 | 5.73% | **5.34%** | 5.66% | **5.28%** |
| RNASim-1000_C_500 | **7.59%** | 7.73% | **7.44%** | 7.59% |

## 4 Conclusions

Multiple sequence alignment is a basic step in many bioinformatics pipelines, with phylogeny estimation one of the applications of interest. Here we have shown that a small change to the PASTA pipeline yields a consistent improvement in alignment accuracy on a collection of biological and simulated datasets. Although the improvement was small, it is suggestive of a trend that could lead to bigger improvements on larger datasets. More generally, it suggests also the possibility of using criteria, such as maximum likelihood scores, to select between competing multiple sequence alignments computed for the same dataset. In general, future work is needed in order to better understand how to estimate multiple sequence alignments.

## Acknowledgments

This work was performed while the first author was an undergraduate, and participating in a Research Experience for Undergraduates program in the Department of Computer Science at the University of Illinois at Urbana-Champaign.

